# POT-1 telomere binding protein promotes a novel form of Transgenerational Epigenetic Inheritance

**DOI:** 10.1101/687988

**Authors:** Evan H. Lister-Shimauchi, Michael Dinh, Paul Maddox, Shawn Ahmed

## Abstract

Transgenerational Epigenetic Inheritance occurs when gametes transmit forms of information without altering genomic DNA^1^. Although deficiency for telomerase in human families causes transgenerational shortening of telomeres^2^, a role for telomeres in Transgenerational Epigenetic Inheritance is unknown. Here we show that Protection Of Telomeres 1 (Pot1) proteins, which interact with single-stranded telomeric DNA^3,4^, function in gametes to regulate developmental expression of telomeric foci for multiple generations. *C. elegans* POT-1 and POT-2^5,6^ formed abundant telomeric foci in adult germ cells that vanished in 1-cell embryos and gradually accumulated during development. *pot-2* mutants displayed abundant POT-1::mCherry foci throughout development. *pot-2* mutant gametes created F1 cross-progeny with constitutively abundant POT-1::mCherry and mNeonGreen::POT-2 foci, which persisted for 6 generations but did not alter telomere length. *pot-1* mutant and *pot-2; pot-1* double mutant gametes gave rise to progeny with constitutively diminished Pot1 foci. Genomic silencing and small RNAs potentiate many transgenerational effects^7^ but did not affect Pot1 foci. We conclude that *C. elegans* POT-1 functions at telomeres of *pot-2* mutant gametes to create constitutively high levels of Pot1 foci in future generations. As regulation of telomeres and Pot1 have been tied to cancer^8,9^, this novel and highly persistent form of Transgenerational Epigenetic Inheritance could be relevant to human health.

---

Telomeres cap the ends of linear chromosomes and are typically composed of simple repetitive sequences that are maintained by *de novo* telomere repeat addition by telomerase^2^. The *C. elegans* Pot1 homologs POT-1 and POT-2 ^10^ repress telomerase activity at telomeres ^6^, and both individual proteins can bind to the base of T-loops, which form when telomeric single-stranded overhangs fold back to create strand-invasion intermediates ^5,11^. POT-2 but not POT-1 represses the telomerase-independent telomere maintenance process termed Alternative Lengthening of Telomeres (ALT) ^12^. Therefore, these Pot1 proteins have both common and distinct telomeric functions.

A single-copy transgene that expresses POT-1::mCherry was previously shown to result in nuclear mCherry foci at telomeres of adult *C. elegans* germ cells and mature sperm ^6^. We found that POT-1::mCherry foci became weaker as oocytes matured (Fig. S1A,B) and that fertilization of oocytes resulted in 1-cell embryos that lacked POT-1 foci in both parental pronuclei and also in the interphase zygotic nucleus (Fig. 1A-D). The number of POT-1 foci gradually increased during embryonic development to 8.0+/−0.5 foci per nuclei in 32-cell embryos (Fig. 1I). We were able to quantify POT-1::mCherry foci in a small fraction of germ cells at L1 to L4 larval stages (Fig. S1E-N) and found that an abrupt transition to the previously observed ~19 POT-1::mCherry foci per adult mitotic germ cell nucleus ^6^ occurred as germ cells of L4 larvae matured into adult germ cells (Fig. S1O). This indicates that Pot1 foci go through two dramatic transitions during development.

**Figure 1.**
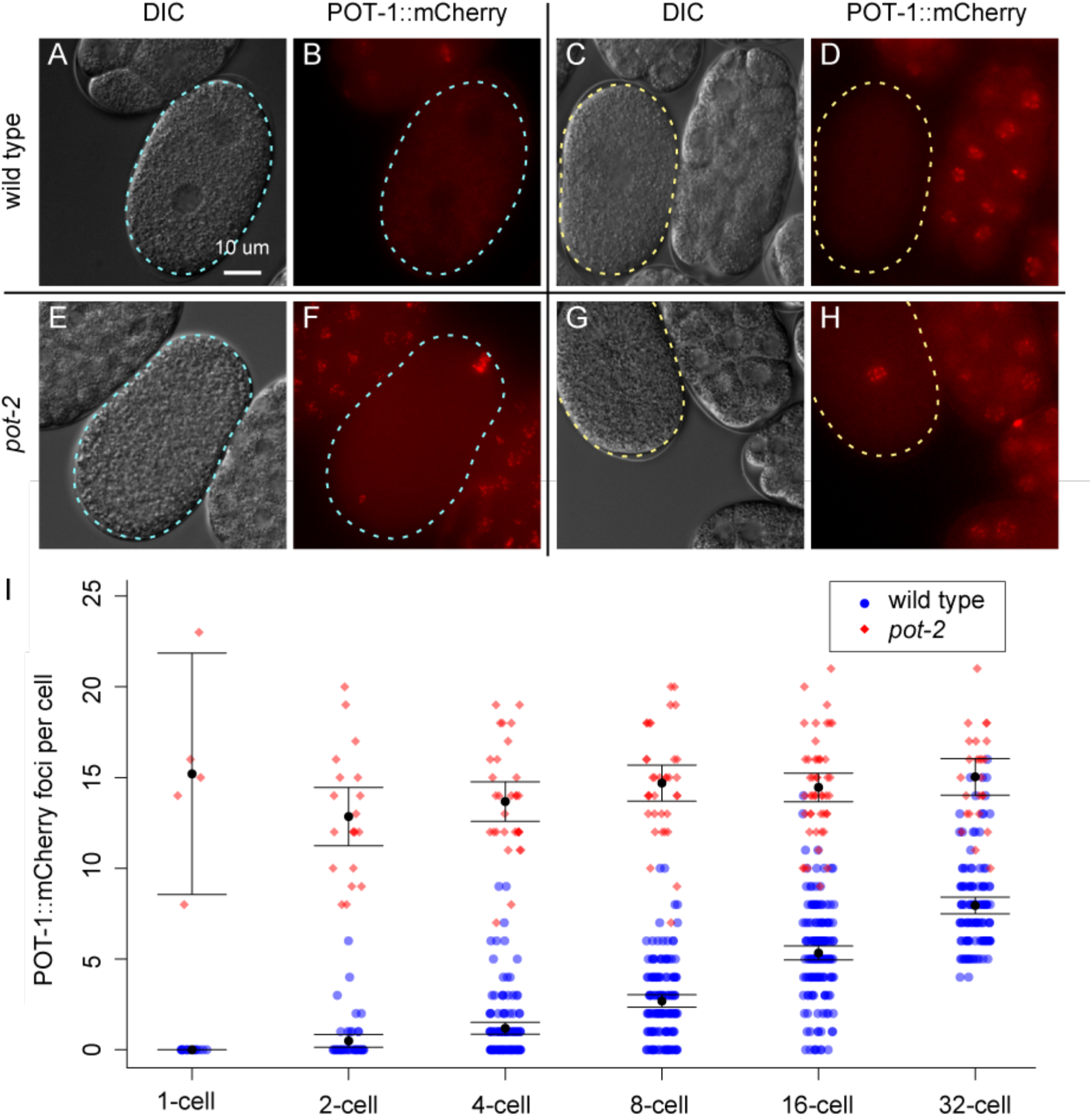
Pot1 foci number increases during early embryonic development. A-H) DIC and fluorescent images of a 1-cell embryos before (A,B,E,F) and after (C,D,G,H) pronuclei fusion in either a wild-type (A-D) or *pot-2* mutant (E-H) background. Cyan and yellow dashed lines indicate 1-cell embryos before and after pronuclei fusion, respectively. I) POT-1::mCherry foci in individual cells of embryos from 1-to 32-cell stage in wild type (blue circles) and *pot-2* mutants (red diamonds). The difference between wild type and *pot-2* mutants is significant at every time point (Wilcox p < 10^−5^).

Deficiency for *pot-2* was previously shown not to affect the abundance of POT-1::mCherry foci in meiotic cells of the adult germline ^6^. However, we found that the three most proximal (mature) oocytes of *pot-2* mutants had many POT-1::mCherry foci (11.4+/−0.8 foci per nucleus) (Fig. S1C,D). Abundant POT-1::mCherry foci were also observed in pronuclei and interphase nuclei of 1-cell *pot-2* mutant embryos (15.2 +/− 4.7 foci per nucleus) (Fig 1E-H), which contrasts sharply with the absence of POT-1 foci in wild-type control 1-cell embryos (Fig. 1A-D). Mutation of *pot-2* causes gradual telomere elongation over the course of 20 generations ^6^, and the *pot-2* mutant strain expressing POT-1::mCherry in 1-cell embryos had been cultured for more than 50 generations prior to imaging. We therefore examined POT-1::mCherry foci using a *pot-2(tm1400)* mutation that was outcrossed 14 times to create a strain whose telomere length is close to wild type ^6^ (Fig. S1P). We observed high levels of POT-1::mCherry foci in all 1- and 2-cell embryos derived from progeny of pot-2 −/− animals whose mothers were *pot-2* +/− heterozygotes. We also found that heterozygosity for *pot-2* does not result in abundant POT-1::mCherry foci in early embryos. Therefore, loss of POT-2, rather than long telomeres, results in the immediate appearance of abundant POT-1::mCherry foci in the interphase nuclei of early embryos.

Given the strong effect of deficiency for *pot-2* on early embryonic POT-1::mCherry foci, we introduced an *mNeonGreen* tag at the endogenous *pot-2* locus using CRISPR/Cas9-mediated genome modification ^13^. A cytoplasmic pool of mNeonGreen::POT-2 was present at all embryonic stages and in all adult germ cells (Fig. S2A-T). Like POT-1::mCherry, mNeonGreen::POT-2 formed very strong nuclear foci in mature sperm (Fig. S2M-P), and embryonic mNeonGreen::POT-2 nuclear foci colocalized perfectly with POT-1::mCherry foci (Fig. S2Q-T). We found that mNeonGreen::POT-2 foci were absent in late oocyctes and 1-cell interphase embryos and slowly reappeared during embryo development, as observed for POT-1::mCherry (Fig. S2Q-T). Cytoplasmic mNeonGreen::POT-2 was not perturbed at these stages of development, indicating that the loss of nuclear POT-2 foci in wild-type control strains is not a consequence of transcriptional or translational silencing of POT-2.

We asked if deficiency for *pot-2* in either oocytes or sperm would create a parent-of-origin effect that would alter levels of Pot1 foci. When males expressing POT-1::mCherry and mNeonGreen::POT-2 were crossed with *pot-2* mutant hermaphrodites, abundant POT-1::mCherry foci were observed in 1- and 2-cell embryos of *pot-2* heterozygous F1 cross progeny. This phenotype persisted for six generations, even for F2 lines whose progeny all possessed the *mNeonGreen::pot-2* transgene, gradually reverting to wild-type levels of POT-1::mCherry foci by generation F7 (Fig. 2A).

**Figure 2.**
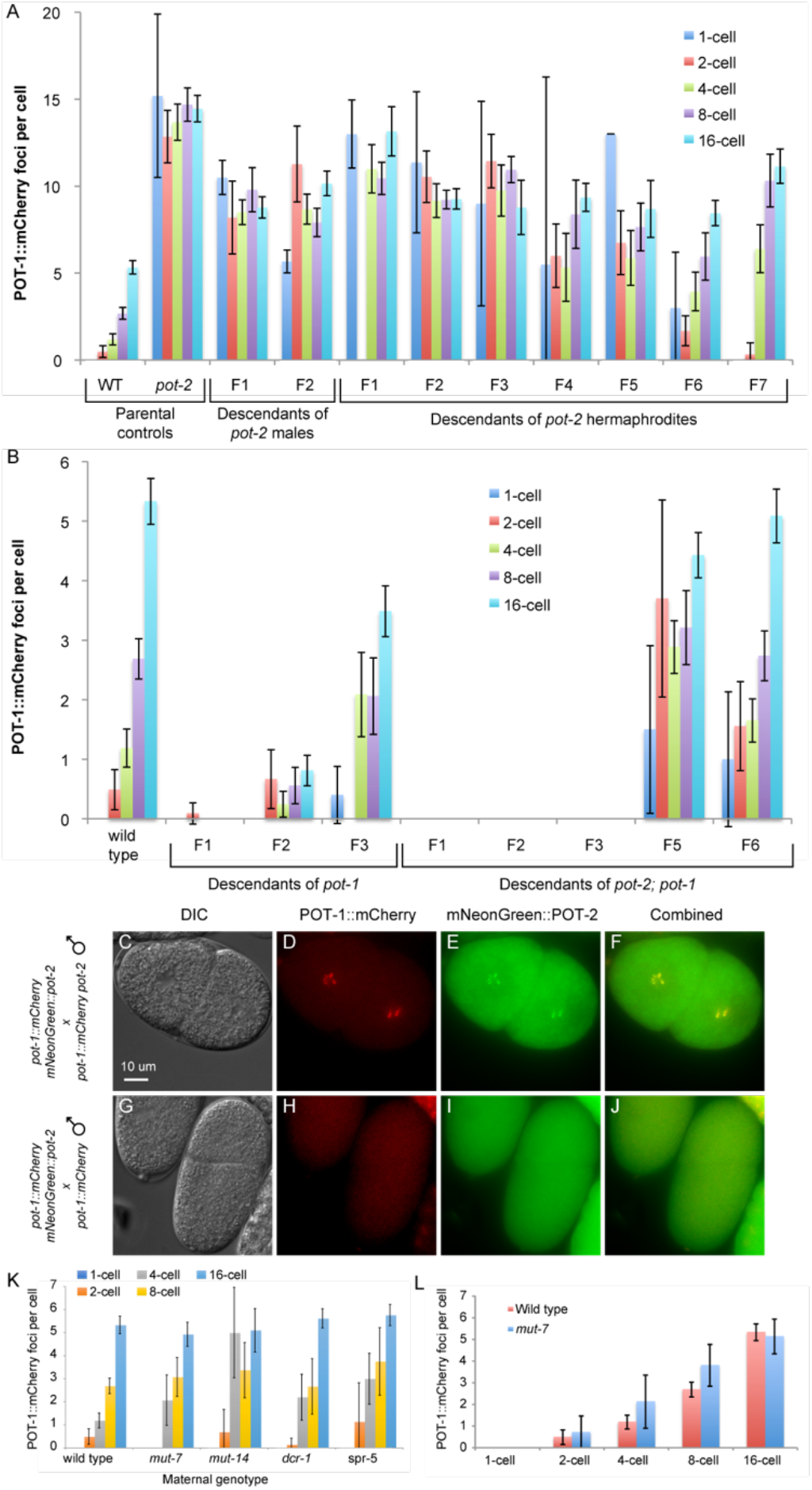
*pot* gene mutations affect Pot1 foci in subsequent generations. A) POT-1::mCherry foci per cell in embryos from P0 parental controls and descendants of either pot-2 mutant males or *pot-2* mutant hermaphrodites crossed to POT-1::mCherry worms. Bar color indicates stage of embryo. B) POT-1::mCherry foci counts per cell in embryos from wild type parental controls and descendants of either *pot-1* or *pot-2; pot-1* mutant males crossed to POT-1::mCherry worms. C-J) F1 2-cell embryos from crosses between *pot-1::mCherry mNeonGreen::pot-2* hermaphrodites and either *pot-1::mCherry pot-2* males (C-F) or *pot-1::mCherry* males (G-J). K) Foci counts in wild-type controls and the descendants of POT-1::mCherry males crossed to hermaphrodites of the indicated genotypes. Compared to wild type using 2-way ANOVA, only *spr-5* is significantly different overall (p = 0.03), although Pot1 foci are absent from 1-cell embryos. L) Foci counts in *mut-7; pot-1::mCherry* homozygous embryos. The difference between wild type and *mut-7* is not significant by Wilcox test at any timepoint.

We asked if the induction of POT-1::mCherry foci in the progeny of *pot-2* mutant hermaphrodites was a maternal effect or a more general form of nuclear inheritance by crossing *pot-2* mutant males with *pot-1::mCherry* hermaphrodites. This resulted in F1 cross-progeny with abundant POT-1::mCherry foci in all early-stage embryos, and this phenotype persisted for at least 2 additional generations (Fig. 2A). An independent cross of *pot-2* mutant males expressing POT-1::mCherry with hermaphrodites expressing POT-1::mCherry and mNeonGreen::POT-2 generated F1 cross-progeny with abundant POT-1::mCherry foci that colocalized with mNeonGreen::POT-2 foci throughout embryonic development, including early embryos (Fig. 2C-F). In contrast, control crosses of *pot-1::mCherry* males and *pot-1::mCherry mNeonGreen::pot-2* hermaphrodites yielded F1 cross-progeny early embryos with no visible Pot1 foci (Fig. 2G-J), and wild-type levels of Pot1 foci were observed at later stages of development. Together, these results indicate gametes that are deficient for *pot-2* transmit a persistent form of nuclear inheritance that results in abundant telomeric foci composed of POT-1 and, somewhat paradoxically, POT-2 in early embryos for multiple generations.

We asked if deficiency for *pot-1* in gametes might affect levels of Pot1 foci by crossing males expressing POT-1::mCherry and mNeonGreen::POT-2 with *pot-1* mutant hermaphrodites. We found that the number of POT-1::mCherry foci per nucleus was either absent or reduced at all stages of embryonic development examined for several generations (Fig. 2B). We also generated an *mNeonGreen::pot-2* strain that was homozygous mutant for *pot-1* and found that mNeonGreen::POT-2 foci were absent from embryos and from most adult germ cell nuclei of this strain, although cytoplasmic mNeonGreen::POT-2 was clearly present (Fig. S2U-X). As deficiency for *pot-1* or *pot-2* in gametes had opposite effects on Pot1 foci in the resulting progeny, we crossed *pot-1::mCherry mNeonGreen::pot-2* males to *pot-2; pot-1* double mutants, singled many F2 animals, examined F2 lines that possessed wild-type *pot-1* and *pot-2* alleles. We found that F1, F2, and F3 embryos resembled those of *pot-1* mutants, with few foci appearing during development, although cytoplasmic mNeonGreen::POT-2 was unaffected in the progeny of *pot-2; pot-1* double mutant gametes (Fig. 2B). Pot1 foci returned to wild-type levels of expression by the F5 generation.

Most examples of Transgenerational Epigenetic Inheritance involve gametes that transmit altered levels of cytosine methylation, histone modifications, or small RNAs that modify gene expression in response to exogenous or endogenous cues ^7,14–16^. Moreover, telomeres in several species including *C. elegans* possess heterochromatic histone modifications such as methylated histone H3K9 ^17^, which can be deposited in response to cues from small RNAs that are homologous to genomic loci. We therefore asked if gametes defective for heterochromatin or for small RNA-mediated genomic silencing would affect Pot1 foci. However, we found that gametes deficient for a gene that promotes genomic silencing, the H3K4 histone demethylase *spr-5* ^18^, or deficient for the small RNA-mediated genomic silencing genes *mut-7, mut-14*, or *dcr-1* ^19^ yielded F1 cross-progeny with F2 embryos in which Pot1 foci were absent at the 1-cell stage and gradually accumulated during embryonic development, as observed for wild type (Fig. 3K). We also generated a *pot-1::mCherry* strain that was homozygous mutant for *mut-7*, which results in transposon activation and abolishes small RNA-mediated heterochromatin^19^, and observed wild-type levels of POT-1 foci in early embryos (Fig. 3L). These data indicate that disrupting small RNAs or H3K9-mediated genomic silencing does not perturb levels of Pot1 foci. We did not test cytosine methylation, as this is absent from *C. elegans*^20^.

**Figure 3.**
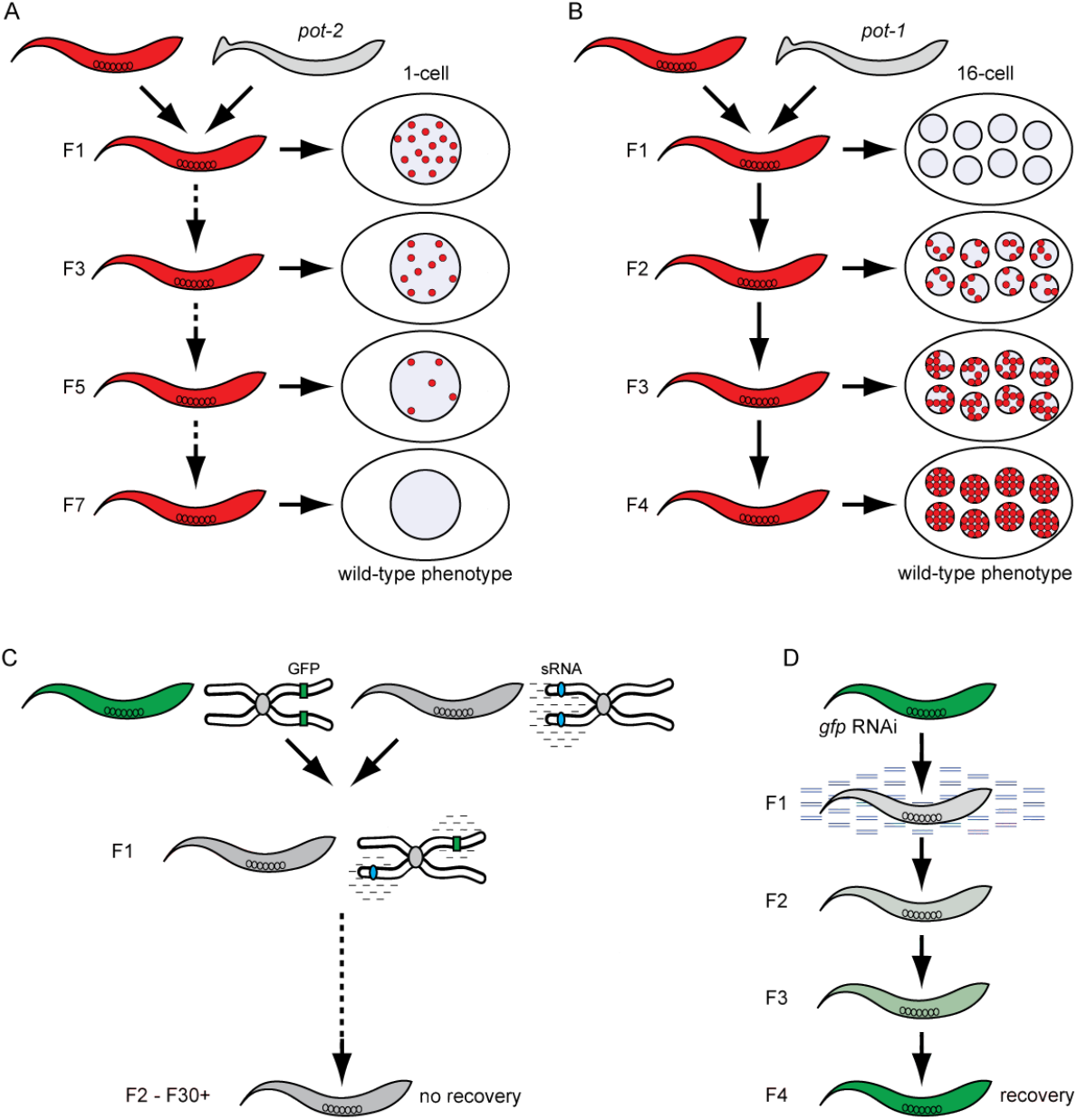
Models of Transgenerational Epigenetic Inheritance. Epigenetic inheritance of Pot1 foci in progeny of *pot-2* (A) and *pot-1* (B) mutant gametes. For comparison, permanent transgenerational silencing via Paramutation (C) and transient transgenerational silencing via RNAi Inheritance (D).

Deficiency for *pot-2* promotes the telomerase-independent telomere maintenance mechanism ALT ^6^, which is active in ~10% of all tumors ^21^. We asked if ALT affects Pot1 foci by crossing *pot-1::mCherry mNeonGreen::pot-2* worms to *trt-1* mutants that had been passaged for over 100 generations under crowded conditions in order to activate the ALT pathway ^19^. We found that F1 cross progeny possessed F2 embryos with wild-type levels of POT-1 and POT-2 foci (Fig. S3A-D), indicting that activation of the ALT pathway does not create gametes whose progeny possess perturbed levels of Pot1 foci.

As *pot-1* or *pot-2* mutant gametes give rise to cross-progeny where levels of Pot1 foci are altered for multiple generations, we asked if telomere length was perturbed in progeny of *pot-1* or *pot-2* mutant gametes. We crossed wild-type males to well-outcrossed early generation *pot-1* or *pot-2* mutant hermaphrodites that possessed telomeres that were modestly longer than wild type, or with control wild-type hermaphrodites. We identified F2 cross progeny that were wild-type for *pot-1* or *pot-2* by PCR. DNA from the descendents of these genetically wild-type F2 was analyzed by Southern blot using a probe for telomeric repeats. We found that well-outcrossed *pot-1* or *pot-2* mutant lines had telomeres that were modestly elongated ^6^, but when either strain was crossed with wild type and genetically wild-type F2s were isolated, descendants of either *pot-1* or *pot-2* mutant gametes displayed telomeres that gradually shortened over the course of 4 generations (Fig. S3E,F).

The stability of cytoplasmic POT-2 in conjunction with the absence of POT-2 foci in wild-type 1-cell embryos demonstrated that Pot1 foci are normally dismantled after fertilization (Fig. S3G-I), and we found that the presence of POT-2 in both parental gametes is required to dismantle Pot1 foci. *pot-1* or *pot-2* mutant gametes constitutively reduced or elevated, respectively, the levels of Pot-1 foci for multiple generations, while telomere length was not markedly altered in their progeny. Therefore, although telomerase dysfunction in humans or mice causes transgenerational effects associated with inheritance of telomeres of altered lengths, we have identified a novel form of Transgenerational Epigenetic Inheritance that modulates the developmental expression of Pot1 foci at telomeres in a manner that is both independent of telomere length and inconsequential to telomere length. Loss of POT-2 in either parental gamete resulted in an inability to dismantle embryonic Pot1 foci in a manner that is remembered for six generations, even in the presence of wild-type POT-2, representing one of the most persistent yet transient forms of Transgenerational Epigenetic Inheritance ever documented ^7^.

The ability of telomeres of *pot-1* and *pot-2* mutant gametes to transform developmental expression of Pot1 foci when crossed with wild-type gametes resembles Paramutation, where small RNAs produced from a silent locus can act in trans to induce permanent heritable silencing of a homologous gene (Fig. 3A-D)^22^. A related silencing process, termed RNAi inheritance, occurs when exogenous small RNAs induce transient transgenerational silencing of GFP transgenes ^14,15^. Telomeres can promote an epigenetic effect termed the Telomere Position Effect, which is a form of Position Effect Variegation where genes adjacent to telomeres can be epigenetically silenced in a manner that is mitotically heritable and reversible ^23,24^. However, both Paramutation and Position Effect Variegation are genomic silencing processes, and we found that defects in heterochromatin formation or small RNA-mediated genome silencing did not affect the developmental expression of Pot1 foci in embryos.

We addressed the remarkable memory of descendants of *pot-2* mutant gametes by studying the progeny of *pot-1; pot-2* double mutant gametes, which did not display Pot1 foci. This implies the presence of POT-1 at telomeres of *pot-2* mutant gametes is an integral component of the memory that modulates this novel form of Transgenerational Epigenetic Inheritance. Jean Baptiste Lamarck and Charles Darwin suggested that it might be beneficial if metazoans could promote the fitness of future generations in response to environmental stimuli ^25^. In this regard, previous analysis of the genomes of 152 wild *C. elegans* strains revealed that 12 possessed a putative inactivating mutation in *pot-2* that was associated with long telomeres, suggesting that either POT-2 or telomere length could contribute to fitness in the wild ^26^.

We conclude that Pot1 proteins regulate a novel and persistent form of Transgenerational Epigenetic Inheritance that concerns developmental expression of Pot1 foci at telomeres. Regulation of telomeres has been tied to aging and cancer, and mutations of the Pot1 locus in human families are associated with an increase in the incidence of cancer ^8,9^. If Transgenerational Epigenetic Inheritance of Pot1 foci were relevant to human health, then disease might be observed in human pedigrees segregating Pot1 mutations for family members that lack the Pot1 mutation itself ^9^.

## Methods

### Strains

All strains were cultured at 20°C on Nematode Growth Medium (NGM) plates seeded with *E. coli* OP50. Strains used include Bristol N2 wild type, *ypSi2 (pdaz-1::pot-1::mCherry::ttb-2 3’UTR), trt-1(ok410) I, spr-5(by101) I, spr-5(by134) I, pot-2(tm1400) II, pot-2(ypSi3 [mNeonGreen::pot-2]) II, unc-52(e444) II, pot-1(tm1620) III, mut-7(pk720) III, dcr-1(mg375) III*, and *mut-14(pk738) V*.

### Generation of fresh pot-2 homozygous mutants from balanced stocks

The *pot-2* locus and the *pot-1::mCherry* transgene both reside on chromosome II. *pot-2* is very close to *unc-52*, while *pot-1::mCherry* is very close to *rol-6*. We created heterozygous *pot-2(tm1200) − / +* males by balancing with an *unc-52* mutant allele. We crossed these with *pot-1::mCherry* hermaphrodites to create *pot-2* − / + F1 hermaphrodites possessing a single copy of *pot-1::mCherry* (Fig. S1P). *pot-2* heterozygotes possessing two copies of *pot-1::mCherry* were obtained by selecting F2 worms that produced some *unc* progeny and only progeny that possessed *pot-1::mCherry*. Most F3 embryos produced by these F2 worms did not display abundant early foci. However, a few, most likely the *pot-2* homozygous mutants, did display early foci. We also studied F3 adult progeny that were *pot-1::mCherry; pot-2 − / −* homozygotes and found high levels of POT-1::mCherry foci in all 1- and 2-cell F4 embryos. *pot-2* mutants that are created from well-outcrossed *pot-2* mutant stocks have normal telomere lengths at generation F4, which slowly increase in length over the course of 16 generations ^6^.

### Generation of *mNeonGreen::pot-2 transgenic strain*

DNA constructs for the *mNeonGreen::pot-2* transgene were generated based on the previously described SapTrap protocol^27^. CRISPR-based insertion into the genome was accomplished using previously described protocols ^13^. Sanger sequencing of the mNeonGreen tag and neighboring genome was used to confirm correct insertion.

We generated endogenous *pot-1* tags through similar means. Though we were successful in generating both N- and C-terminal mKate tags of *pot-1*, we were unable to visualize the transgenes. This may be due to weak expression of the endogenous *pot-1* locus. Expression of *pot-2* is considerably higher than *pot-1* based on RNAseq data from the modENCODE project ^28^. We would therefore expect the signal from an endogenous *pot-1* transgene to be even weaker than the *mNeonGreen::pot-2* trasngene.

### Crosses

To perform crosses, 24 males and 10 hermaphrodites were placed on a single plate. Plates were placed at 15°C for 3 days to obtain F1 cross-progeny embryos.

### Imaging

Germ cells and embryos were mounted on 2% agarose pads and imaged live using a DeltaVision Elite microscope (Applied Precision) with a 60x Silicon oil objective lens. For germline imaging, immobilization was assisted by transferring worms into 5uL drops of 0.2 mM levamisole on agarose pads. For embryo imaging, adults were extruded in M9 buffer and embryos were transferred onto agarose pads.

### Quantification and statistics

Images were analyzed using ImageJ software. Foci were counted manually from fluorescent images, while embryo stage was determined from DIC images.

Statistical calculations were performed using R software. Foci numbers were compared using either the Wilcoxon rank-sum test or 2-way ANOVA, depending on the number of variables tested simultaneously. When making multiple simultaneous statistical comparisons, p-values were adjusted using the Holm-Bonferroni method. All error bars displayed represent 95% confidence intervals.

## Acknowledgements

Felix Peng contributed to the initial observations of the repression of POT-1::mCherry foci in early embryos by POT-2. Dan Dickinson provided plasmids used in the generation of the *mNeonGreen::pot-2* targetting construct. We thank Bob Goldstein and Dan Dickinson for advice regarding CRISPR-mediated genome modifications. We thank Amy Maddox for training on and access to her microscope. This work was supported by NIH R01 066228 to SA.

## Author Contributions

E.H.L.-S., M.D., P.M. and S.A. designed the experiments. E.H.L.-S., M.D. and S.A. carried out the experiments, E.H.L.-S. and S.A. wrote the manuscript.

---

**Supplemental Figure 1.**
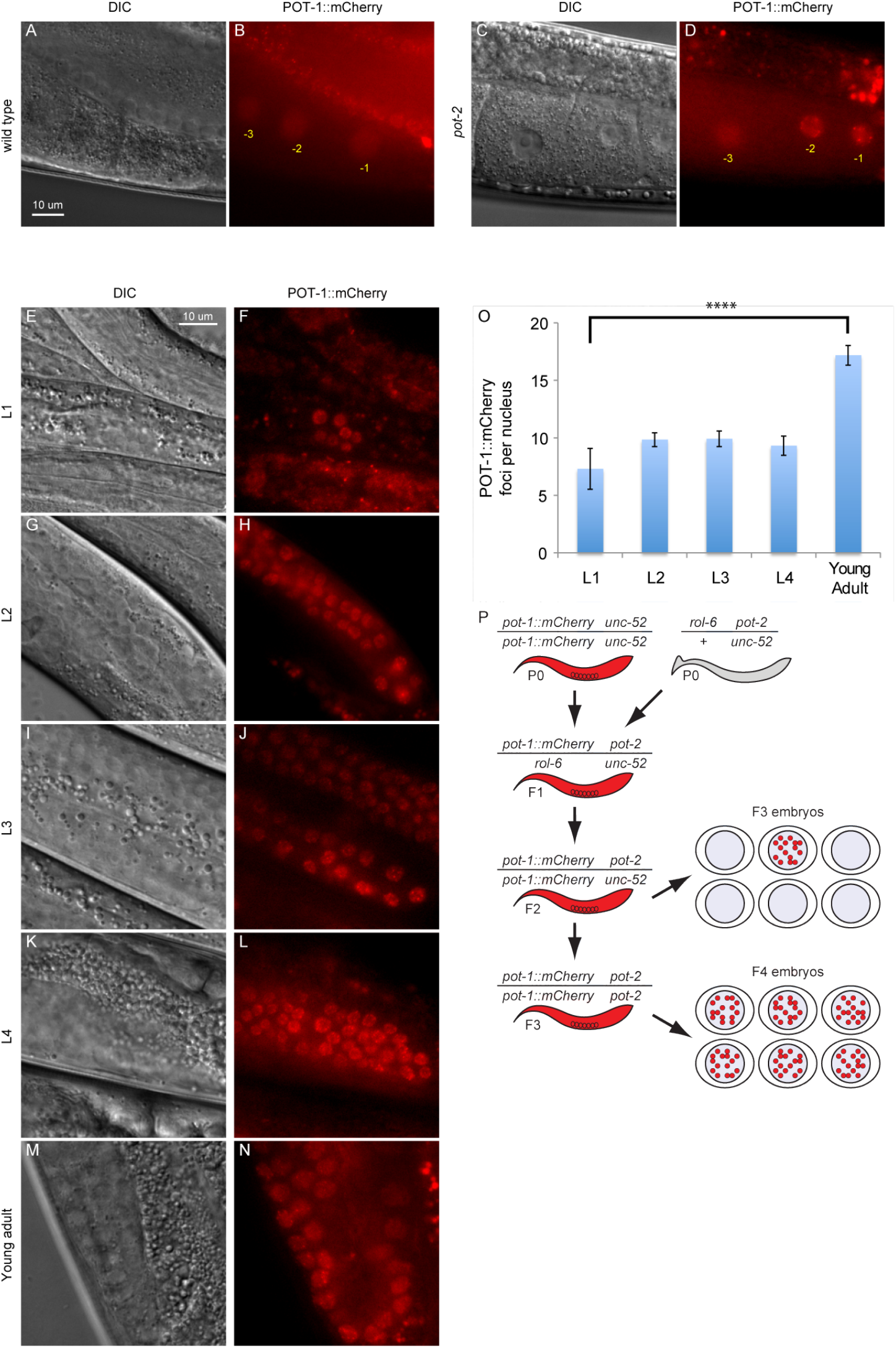
A-D) POT-1::mCherry foci in a wild type (A, B) and *pot-2* mutant (C-D) background. Oocyte position is indicated by yellow text. E-N) POT-1::mCherry in the distal region of L1-L4 larvae (E-L) and young adult worms (M, N). O) Quantification of POT-1::mCherry in mitotic zone cells of L1-L4 larvae and young adults. The only stage significantly different from L1 is young adult (Wilcox p<10^−5^). P) Diagram of crosses using outcrossed *pot-2* mutants with wild-type telomere lengths.

**Supplemental Figure 2.**
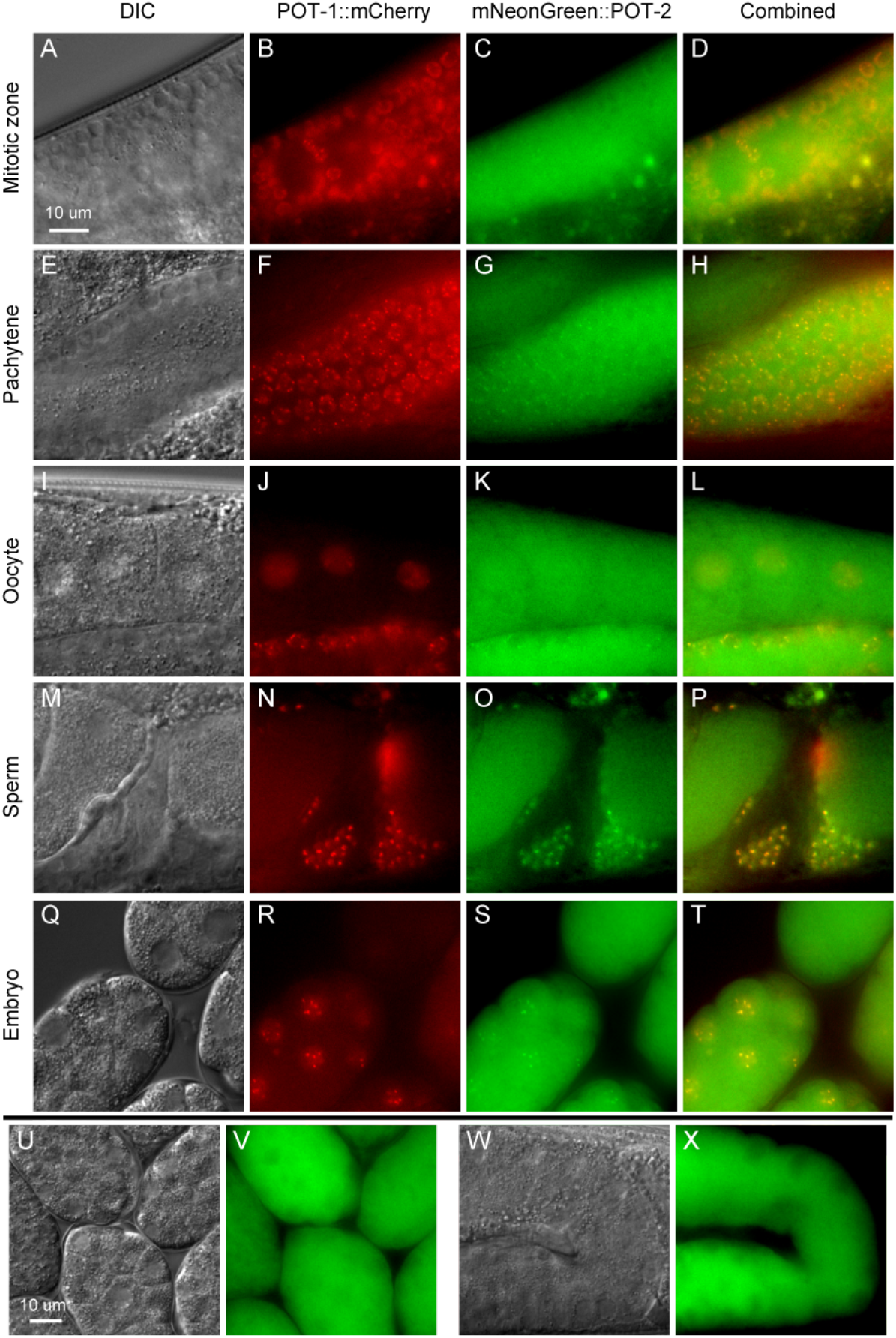
A-T) DIC and fluorescent images of *pot-1::mCherry mNeonGreen::pot-2* worms. Representative images from the germline mitotic zone (A-D), pachytene zone (E-H), oocytes (I-L), sperm (M-P), and embryos of early and late stages (Q-T). U-X) *mNeonGreen::POT-2* localization in the embryos (U, V) and germline (W, X) of *pot-1* mutant homozygotes.

**Supplemental Figure 3.**
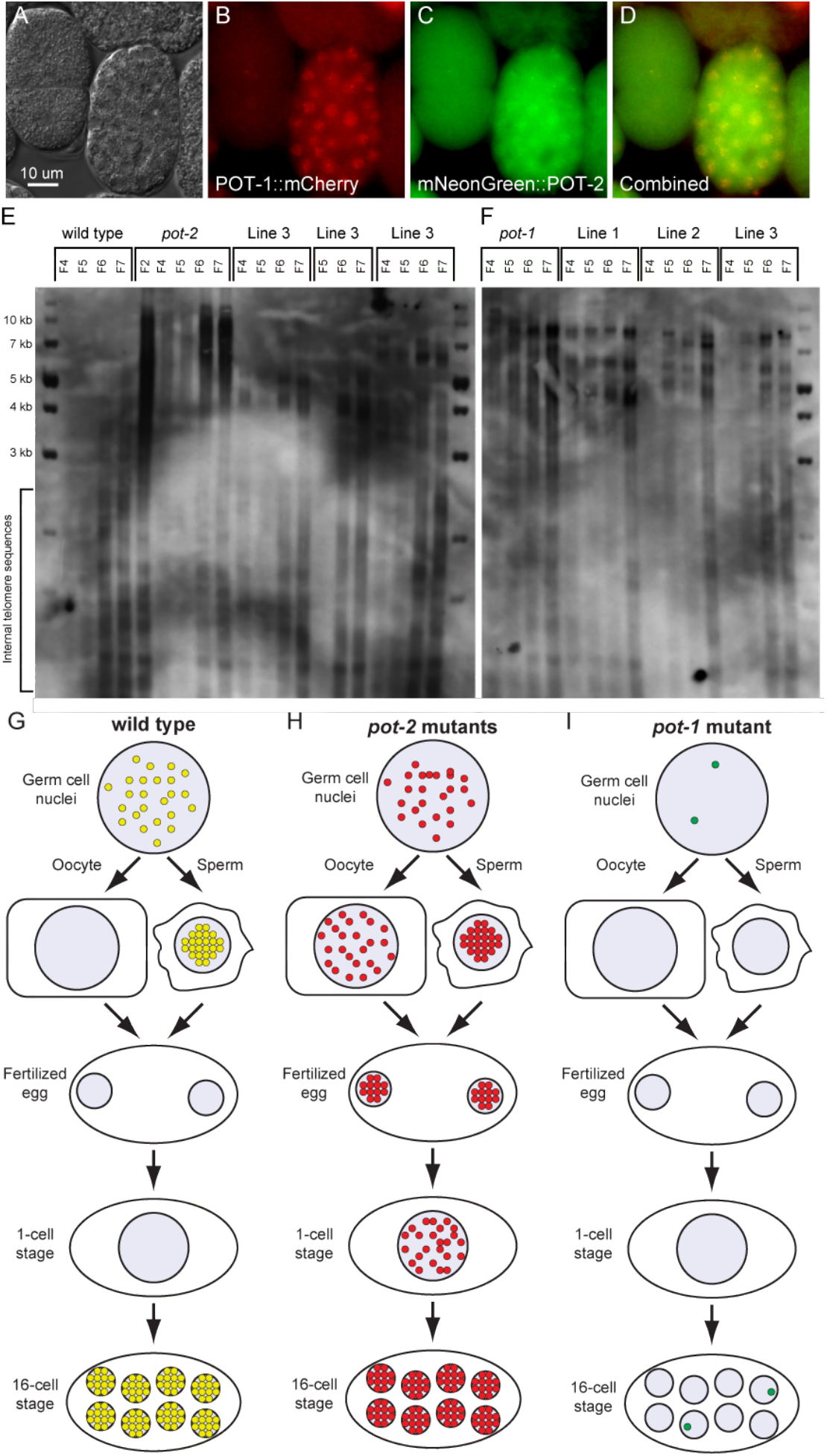
A-D) POT-1::mCherry and mNeonGreen::POT-2 expression in *trt-1* homozygous mutants propagated by ALT. E,F) Southern blot using DNA from the genetically wild-type descendents of crosses between wild type and *pot-2* (E) or *pot-1* (F) mutant worms. G-I) Descriptive model of the developmental timing of POT foci localization in wild type (G), *pot-2* mutants (H), and *pot-1* mutants (I).

